# Maintenance of pyrophosphate homeostasis in multiple subcellular compartments is essential in *Plasmodium falciparum*

**DOI:** 10.1101/2025.02.20.639246

**Authors:** Ikechukwu Nwankwo, Hangjun Ke

## Abstract

Pyrophosphate is a byproduct of numerous cellular reactions that use ATP or other nucleoside triphosphates to synthesize DNA, RNA, protein, and other molecules. Its degradation into monophosphate is thus crucial for the survival and proliferation of all life forms. The human malaria parasite *Plasmodium falciparum* encodes two classes of pyrophosphatases to hydrolyze pyrophosphate. The first consists of *P. falciparum* proton pumping vacuolar pyrophosphatases (PfVP1 and PfVP2), which localize to the parasite’s subcellular membranes and work as proton pumps. The second includes *P. falciparum* soluble pyrophosphatases (PfsPPases), which have not been well characterized. Interestingly, the gene locus of PfsPPase encodes two isoforms, PfsPPase1 (PF3D7_0316300.1) and PfsPPase2 (PF3D7_0316300.2). PfsPPase2 contains a 51- amino acid organellar localization peptide that is absent in PfsPPase1. Here, we combine reverse genetics and biochemical approaches to identify the localization of PfsPPase1 and PfsPPase2 and elucidate their individual functions. We show that PfsPPases are essential for the asexual blood stage. While PfsPPase1 solely localizes to the cytoplasm, PfsPPase2 exhibits multiple localizations including the mitochondrion, the apicoplast, and, to a lesser extent, the cytoplasm. Our data suggest that *P. falciparum* has taken a unique evolutionary trajectory in pyrophosphate metabolism by utilizing a leader sequence to direct sPPases to the mitochondrion and apicoplast. This differs from model eukaryotes as they generally encode multiple sPPases at distinct genetic loci to facilitate pyrophosphate degradation in cytosolic and organellar compartments. Our study highlights PfsPPases as promising targets for the development of novel antimalarial drugs.

## Introduction

Malaria is a severe infectious disease that threatens nearly half of the world’s population. According to a recent WHO report, about 247 million malaria cases were recorded and ∼ 619,000 lives were killed each year [1]. The disease is caused by protozoans belonging to the *Plasmodium* genus and of five human malaria species, *P. falciparum* is the most virulent, affecting a large population, especially in the tropical and sub-tropical regions of Africa. Despite the efforts put in place to control and eradicate the disease, malaria control progress has been declining recently due to several factors, including the rapid emergence of drug-resistant parasites, which gradually renders most antimalarial drugs ineffective [2]. As a result, it is urgent to continuously investigate the parasite biology to lay the groundwork for developing novel therapeutic treatments.

Inorganic pyrophosphatases (PPases) play a crucial role in modulating the cellular concentrations of pyrophosphate (PPi) [3], a metabolic byproduct of numerous cellular reactions involving the use of ATP or other nucleoside triphosphates in the synthesis of DNA, RNA, proteins, and other molecules [4]. They catalyze the hydrolysis of PPi into two molecules of monophosphate (Pi) through an irreversible reaction, which has been identified to be thermodynamically favorable, resulting in the shift of the overall cellular equilibrium towards biosynthesis [5]. The absence of PPases would result in the accumulation of PPi to toxic levels, which in turn affects numerous biological processes in the cells; therefore, PPases are essential in all organisms. In *C. elegans*, null mutations of its PPase led to developmental arrest at the larval stage as evidenced by defects in the animal’s intestine [6]. Mutations of PPase in the budding yeast (*S. cerevisiae*) also cause cell cycle arrest and cell death [7]. Furthermore, various studies on pathogenic organisms have focused on PPases as potential drug targets. For example, the drug-resistant strains of *Staphylococcus aureus* are highly susceptible to small molecule inhibitors against PPases [8].

Unlike fungi and metazoans that only express soluble pyrophosphatases, *P. falciparum* encodes two distinct kinds of inorganic pyrophosphatases, including the soluble pyrophosphatase (*P. falciparum* soluble pyrophosphatase, PfsPPase) and the membrane-bound H^+^ translocating vacuolar pyrophosphatases (*P. falciparum* vacuolar pyrophosphatases, PfVP1 and PfVP2) [9]. Recently, we have demonstrated that PfVP1 is uniquely localized to the parasite plasma membrane [10], showing a sharp contrast to other parasites such as *Toxoplasma gondii* and Kinetoplastids whose VP1 proteins localize to intracellular organelles called acidocalcisomes [11–14]. We further demonstrated that ring stage parasites utilize PfVP1 to maintain the plasma membrane potential, cytosolic pH, and energy homeostasis, likely enabling them to allocate the limited ATP generated via a low level of glycolysis to other energy-costly processes [10]. On the other hand, the functions of *P. falciparum* soluble pyrophosphatases (PfsPPases) remain not well characterized.

Interestingly, the genetic locus of PfsPPase (PF3D7_0316300), as depicted in PlasmoDB (www.plasmoDB.org), encodes two isoforms: PF3D7_0316300.1 (380 aa) and PF3D7_0316300.2 (431 aa), which we henceforth refer to as PfsPPase1 and PfsPPase2, respectively. Both enzymes have the same peptide sequence except that PfsPPase2 contains a putative organellar localization signal (51-aa) at its N-terminus (**Figure 1A**). The 51-aa leader sequence contains positively charged and hydrophobic amino acid residues (**Figure 1A**), which resemble the features of mitochondrial targeting sequences [15–17]. Further, MitoProt predicts PfsPPase2 to be mitochondrially localized with a high confidence score of 0.8, suggesting that PfsPPases are not only cytosolic but may also be localized to subcellar organelles. However, this possible organellar localization of PfsPPases was not reported in the previous study led by Jamwal *et al.*, as the authors solely focused on one isoform (PfsPPase1) but not two [18].

**Figure 1.**
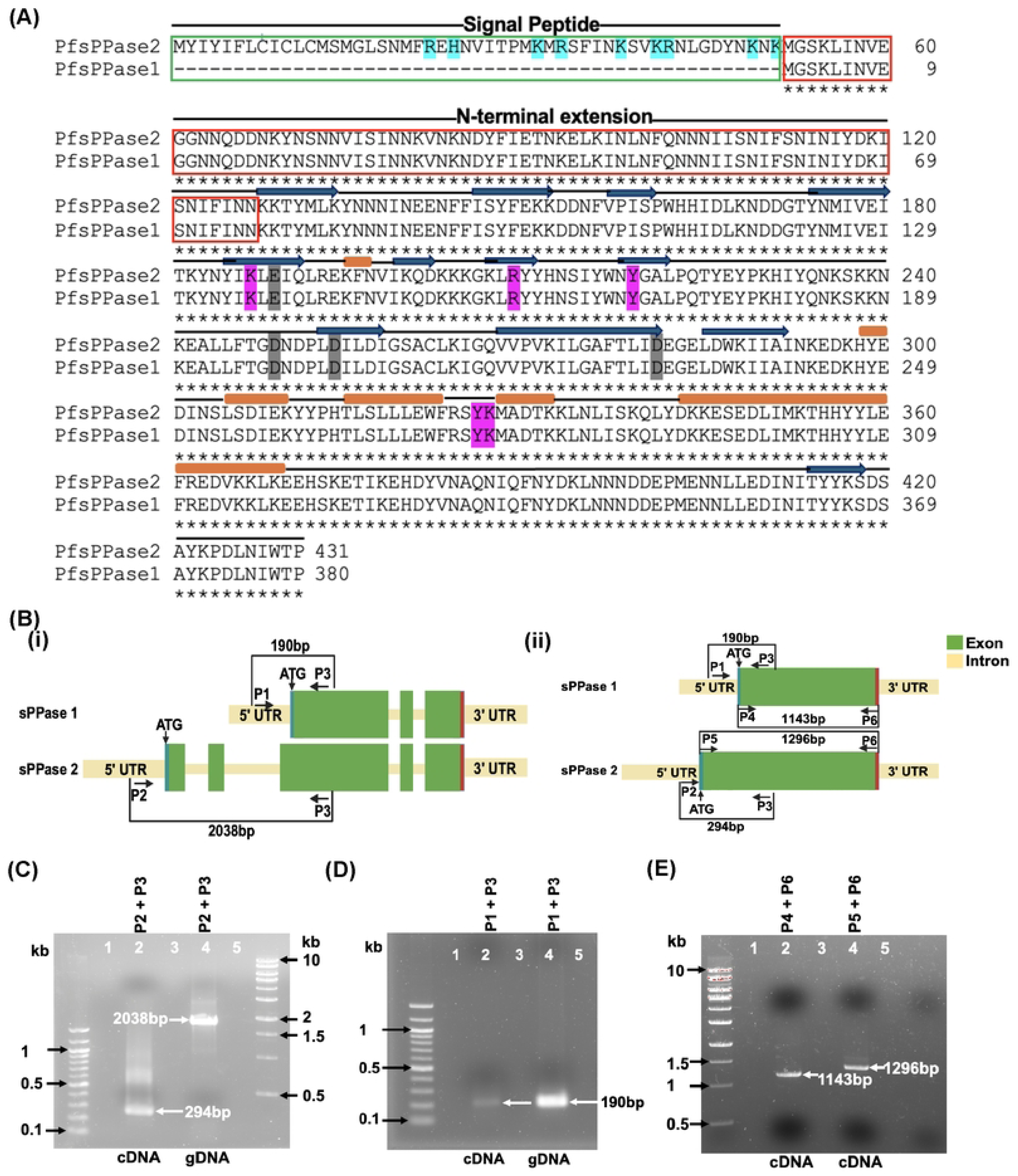
The gene locus of PfsPPases. A, Protein sequence alignment of PfsPPase1 and PfsPPase2. The 51-aa leader sequence in PfsPPase2 is highlighted in a green box. The N-terminal extension of PfsPPases is shown in a red box. PfsPPases are featured with beta strands (dark blue arrows) and alpha helixes (brown rectangular bars). Some critical amino acids are shaded in magenta (involved in pyrophosphate binding) or grey (involved in metal binding). This figure is modified from the previous study [18]. B, Schematic representation of the PfsPPases genetic locus as depicted in PlasmoDB (www.PlasmoDB.org). (i) with introns, (ii) without introns. C, Gel electrophoresis of PfsPPase2 amplified from cDNA and gDNA by primers P2+P3. D, Gel electrophoresis of PfsPPase1 amplified from cDNA and gDNA by primers P1+P3. E, The full-length transcripts of both PfsPPase1 and PfsPPase2 were amplified from cDNA by primers P4+P6 or P5+P6, respectively.

Studies from model eukaryotes suggest that multiple sPPases can be targeted to various subcellular compartments for PPi degradation and homeostasis. For instance, *S. cerevisiae* contains two separate sPPases in the cytosol and the mitochondrion with only 47% identity [19]; *Arabidopsis thaliana* employs six sPPases for PPi degradation in the cytoplasm, the mitochondria, and the plastids [20, 21]. Derived from secondary endosymbiosis, *Plasmodium* parasites possess three genomes within distinct subcellular compartments—the nucleus [22, 23], the mitochondrion [24–26], and the apicoplast (a plastid derivative) [27–30]. Within these compartments, various biological reactions including gene replication/transcription and protein translation generate PPi, which must be hydrolyzed to prevent accumulation of PPi to a toxic level to cause damages. While PPi generated in the nucleus can be trafficked/diffused through the nuclear pores into the cytoplasm and degraded by cytoplasmic sPPases as previously suggested [31], it remains unknown how the mitochondrion and the apicoplast deal with PPi homeostasis in malaria parasites.

In this study, we aim to identify the localization of PfsPPase1 and PfsPPase2 and elucidate their individual functions. Although PfsPPases have previously been predicted to be mutable in *P. falciparum* by a piggyBac mutagenesis study [32], we reveal that both isoforms are essential for *P. falciparum* in the asexual blood stage. Our findings demonstrate that PfsPPase1 localizes to the parasite cytoplasm whereas PfsPPase2 exhibits multiple localizations including the mitochondrion, the apicoplast, and to a lesser extent, the cytoplasm. We conclude that degradation of both cytoplasmic and organellar PPi is essential in maintaining the overall cellular homeostasis of *P. falciparum*.

## RESULTS

### The genetic locus of soluble pyrophosphatase (PfsPPase) in *P. falciparum* encodes two isoforms

The gene locus of PfsPPase (PF3D7_0316300) as observed on PlasmoDB encodes two isoforms: PfsPPase1 (PF3D7_0316300.1) and PfsPPase2 (PF3D7_0316300.2). Their genetic structures are schematically depicted in (**Figure 1B**), with introns present or spliced out.

PfsPPase1 and PfsPPase2 have the same protein sequence except that the latter contains an additional 51-aa leader peptide at the N-terminus (**Figure 1A**), which indicates a possible organellar localization of PfsPPase2. RNA-seq data available on PlasmoDB further supports that the two short exons upstream within the gene locus of PfsPPase2 are transcribed despite their long distance to the next exon (over 1300 bp, base pair) (**Supplementary Figure 1**). Since PfsPPase1 possesses the same protein sequence as PfsPPase2 except the leader peptide, we reasoned that PfsPPase1 is either independently transcribed from the genetic locus or a product of alternative splicing from the PfsPPase2 transcript. To differentiate these two possibilities, we designed primers to amplify several regions of the PfsPPases gene locus using complementary DNA (cDNA) and genomic DNA (gDNA) as the template (**Figure 1B**). PCR confirmed that PfsPPase2 is transcribed as expected (**Figure 1C**) as the sizes of amplicons match the annotated gene model in PlasmoDB. On the other hand, PfsPPase1 is independently transcribed as primers P1 and P3 amplify the same product (190 bp) from both cDNA and gDNA (**Figure 1D**). If PfsPPase1 were a splicing product of PfsPPase2, the 190 bp amplicon would not be amplified from cDNA with primers P1 and P3 because the P1 binding region should be spliced out upon intron removal. Additionally, we verified that the full-length transcripts of both isoforms are present in the parasite cDNA (**Figure 1E**), which display noticeable sizes by DNA gel electrophoresis. Together, these results indicate that PfsPPase1 and PfsPPase2 are independently transcribed from the same genetic locus. This genetic configuration of PfsPPases was not noticed in the previous study that characterized their enzymatic activities [18].

### PfsPPases are essential for parasite development and growth

To directly examine the essentiality of PfsPPases in the parasite, we employed CRISPR/Cas9 [33, 34] and generated a conditional knockdown line in NF54attB [35] using the TetR-DOZI-aptamer system [36, 37] (**Supplementary Figure 2A**). The parasite line was named NF54attB- PfsPPases-3HA^APT^ in which the gene locus of PfsPPases was endogenously tagged with a triple hemagglutinin (3HA) tag at the C-terminus. The genotype of NF54attB-PfsPPases-3HA^APT^ was confirmed by PCR (**Supplementary Figure 2B**). The parasites were cultured in media supplemented with 250 nM of anhydrotetracycline (aTc) to sustain PfsPPases expression (**Supplementary Figure 2C-E**). Next, we tightly synchronized the culture, initiated knockdown studies by aTc removal in Percoll-isolated schizont stage parasites and examined parasite development over several intraerythrocytic developmental cycles (IDCs). After aTc was removed, we confirmed the reduction of the endogenously 3HA-tagged PfsPPase proteins at 48 h and 96 h post knockdown (**Figure 2A**; **Supplementary Figure 3A**). Our C-terminal tagging strategy should theoretically produce two tagged proteins with noticeable differences in molecular weights, 45 kDa (PfsPPase1-3HA, 380 aa plus tag) and 51 kDa (PfsPPase2-3HA,431 aa plus tag). However, western blot only detected one band near the anticipated molecular weight of PfsPPase1 (45 kDa), suggesting that the PfsPPase2’s leader peptide was efficiently removed and no precursor protein of PfsPPase2 (51 kDa) was detectable. This phenomenon is not uncommon in *P. falciparum* as mitochondrially localized proteins often exhibit the processed form with no precursors detectable in western blot [38, 39]. Upon the knockdown of PfsPPases, Giemsa-stained images showed that although PfsPPases depleted parasites successfully finished one IDC, they were completely arrested in the trophozoite stage of the second IDC and failed to develop further into schizonts, resulting in a severe block in parasite replication and subsequent lysis of the parasites by the 3rd IDC (Day 6) (**Figure 2B**). We quantified parasitemia of both the control and knockdown cultures over three IDCs and showed that PfsPPases were essential (**Figure 2C**). We further investigated the PfsPPases knockdown parasites under a fluorescence microscope on day 4 post aTc removal when parasite development was arrested (**Figure 2D**). In agreement with western blot data in **Figure 2A**, the PfsPPase proteins were undetectable at 96 h post aTc removal. While the control parasites had already progressed into the schizont stage with an average of 12 nuclei, the PfsPPases knockdown parasites remained stalled/arrested at the late trophozoite stage (or an early schizont stage), exhibiting only 2 or 3 nuclei (**Figure 2D-E**). To investigate whether the developmental arrest observed upon PfsPPases knockdown is due to the accumulation of PPi, we quantified PPi levels. At 48 h of aTc removal, we observed no significant differences in PPi levels between the control and PfsPPases knockdown parasites. However, at 96 h of knockdown, there was a significant increase in PPi levels in the PfsPPases depleted parasites (**Figure 2F**). Altogether, these findings demonstrate that knockdown of PfsPPases causes severe development arrest, elevated PPi levels, and parasite death, validating the essentiality of PfsPPases for *P. falciparum* in the asexual blood stage.

**Figure 2.**
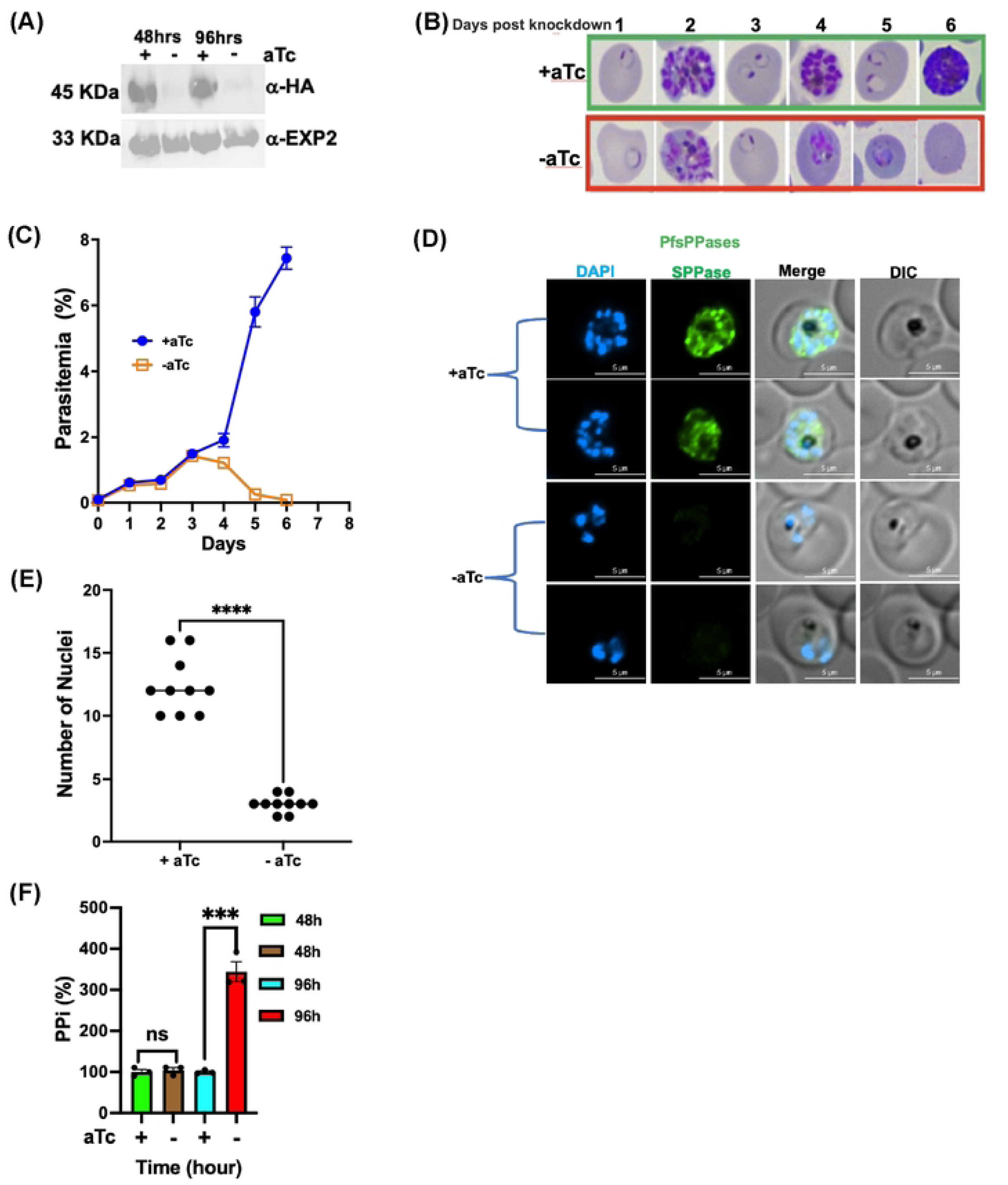
**PfsPPases are crucial for parasite development and growth**. A, Western blot confirming the knockdown of PfsPPases-3HA upon aTc removal in the transgenic parasite line, NF54attB-PfsPPases-3HA^APT^. The blot was probed with anti-HA and re-probed with anti-PfEXP2 to show loading controls. B, Giemsa-stained thin blood smears showing parasite morphological changes after aTc removal for 6 days (3 intraerythrocytic developmental cycles, IDC). Green box, aTc (+). Red box, aTc (-). C, Parasitemia of the knockdown experiment over the knockdown course of 6 days. Parasitemia was determined by light microscopy in triplicate samples. D, Visualization of parasite DNA stained by DAPI post knockdown of PfsPPases for 96 h. Green, PfsPPases-3HA; scale bars, 5 μm. E, Number of nuclei in parasites post knockdown of PfsPPases for 96 h. F, Measurement of PPi in the PfsPPases knockdown parasites after aTc removal for 48 h and 96 h. The concentration of PPi in each condition was measured using the PPi sensor from Sigma and normalized to total protein quantities in each sample, as described previously [10]. The relative PPi concentration of aTc (-) was compared to the aTc (+) control at each timepoint. Statistical analysis was done by t-tests. ***, *p value (*0.006); ns, non-significant. Experiments of A-D have been repeated more than 4 times; panel F was repeated twice.

### Function of two PfsPPase isoforms: partial or full rescue of the PfsPPases knockdown parasites by PfsPPase1 or PfPPase2, respectively

Since our knockdown strategy affects both PfsPPase isoforms simultaneously, the observed phenotypes in **Figure 2** represent the gross defects caused by the loss of both PfsPPase1 and PfsPPase2. To independently characterize the two isoforms and determine their individual functions, we generated two new parasite lines in the available NF54attB-PfsPPases-3HA^APT^. These lines were named as NF54attB-PfsPPases-3HA^APT^-PfsPPase1-3Myc (PfsPPase1-3Myc in short) and NF54attB-PfsPPases-3HA^APT^-PfsPPase2-3Myc (PfsPPase2-3Myc in short), which contain the endogenous PfsPPases tagged with 3HA and aptamers and a constitutively expressed episomal copy of PfsPPase1 or PfsPPase2 tagged with 3Myc. The episomal copy was expressed under the mRL2 (mitoribosomal protein L2) promoter, which has been routinely used to drive protein expression at moderate levels [39, 40]. These two lines, PfsPPase1-3Myc and PfsPPase2-3Myc, allowed us to investigate the function of each isoform after conditional knockdown of the endogenous 3HA-tagged PfsPPases by aTc removal.

In the PfsPPase1-3Myc parasite line, we tightly synchronized the culture, removed aTc, and analyzed the protein levels of the endogenous PfsPPases-3HA and the episomal PfsPPase1-3Myc. Western blot confirmed the constitutive expression of the episomal PfsPPase1-3Myc, while the endogenous 3HA-tagged PfsPPases were efficiently silenced upon aTc removal for 48 h and 96 h, suggesting that parasites solely relied on PfsPPase1-3Myc for PPi hydrolysis in the knockdown conditions (**Figure 3A; Supplementary Figure 3B-C**). To observe the effects of PfsPPase1-3Myc alone on parasite development, we repeated the knockdown experiment and extended the aTc ± cultures for 10 days (5 IDCs). In contrast to the control culture (aTc +), which progressed normally through the ring, trophozoite, and schizont stages of each IDC, we observed that the PfsPPase1-3Myc alone culture (aTc -) started to show growth defects in the 3^rd^ IDC after aTc removal (**Figure 3B**). On day 6, about 40% of the PfsPPase1-3Myc alone culture exhibited abnormal parasites with their morphologies resembling arrested trophozoites undergoing lysis. This abnormal phenotype increased in frequency in the 4^th^ and 5^th^ IDCs post aTc removal, suggesting that a significant portion of parasites was arrested in the trophozoite stage and unable to develop further into the schizont stage. Quantification of parasitemia over the 10 day knockdown course confirmed this developmental arrest (**Figure 3C**). Altogether, these findings show that complementation with PfsPPase1 alone in the PfsPPases knockdown parasites is insufficient to fully rescue the growth and developmental defects observed after depletion of the endogenous PfsPPases, suggesting that parasites need both isoforms of PfsPPases.

**Figure 3.**
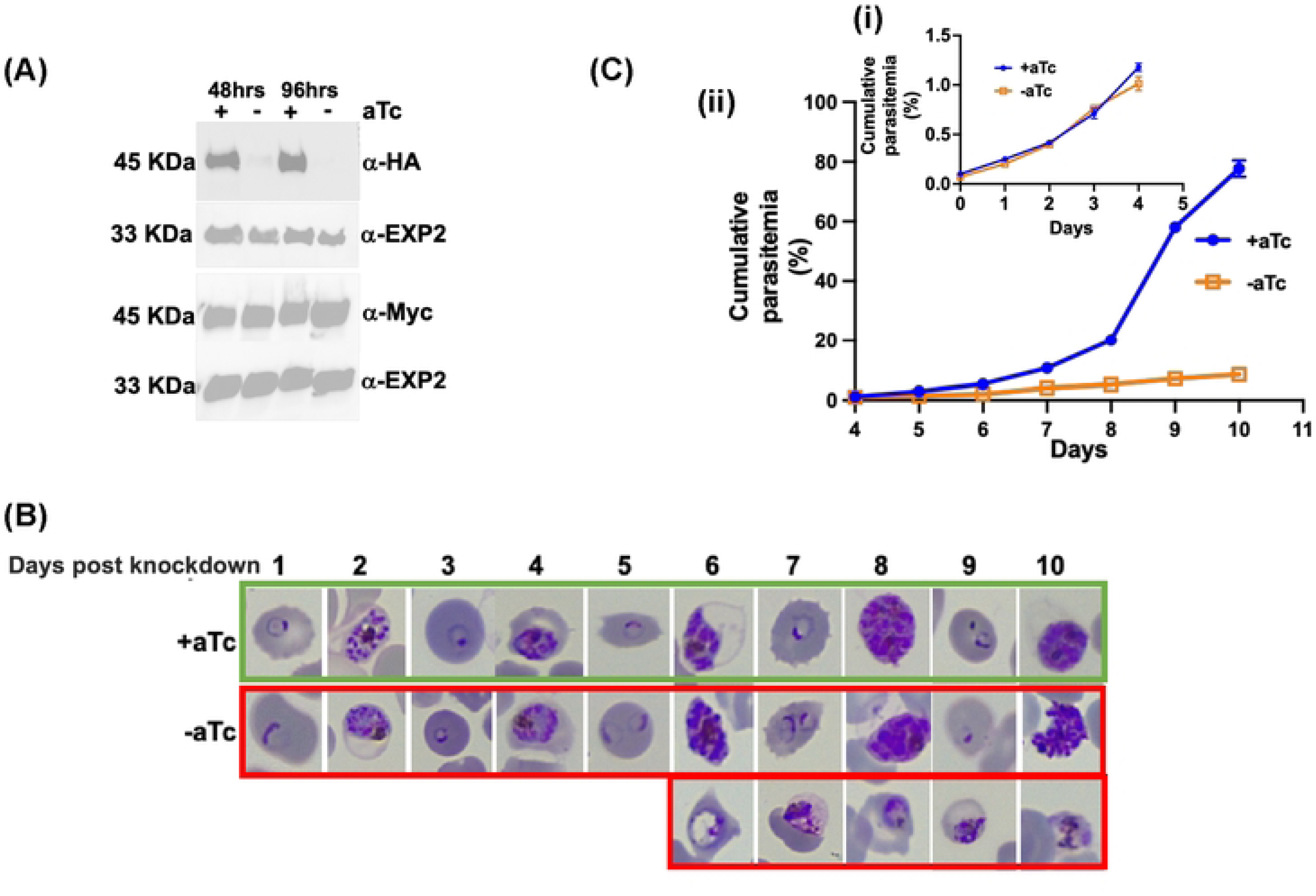
PfsPPase1 partially complements PfsPPases knockdown parasites. A, Western blot analysis of protein lysate from NF54attB-PfsPPase-3HA^APT^-PfsPPase1-3Myc. The blot was probed with anti-HA to verify knockdown of the endogenously tagged PfsPPases-3HA and re-probed with anti-PfEXP2 to show loading controls. Duplicate samples were also blotted with anti-Myc to confirm the expression of the episomally tagged PfsPPase1-3Myc and re-probed with anti-PfEXP2 to show loading controls. B, Giemsa-stained thin blood smears showing parasite morphological changes over a 10-day period after aTc removal. Green box, aTc (+). Red box, aTc (-). C, (i) Parasitemia for the first 4 days of the knockdown experiment. (ii) Parasitemia for the last 6 days. Parasitemia was determined by microscopic counting in at least three biological replicates. These experiments were repeated at least three times.

Using similar approaches, in the PfsPPase2-3Myc parasite line, we assessed the effects of PfsPPase2 alone on parasite development by knocking down the endogenously 3HA-taggged PfsPPases. Again, western blot confirmed that PfsPPase2-3Myc was expressed independently of aTc, while the endogenous 3HA-tagged PfsPPases were effectively knocked down upon aTc removal for 48 h or 96 h (**Figure 4A**; **Supplementary Figure 3D-E**). In contrast to PfsPPase1 that insufficiently rescued the loss of endogenous PfsPPases, PfsPPase2 alone completely sustained parasite development without noticeable defects in parasite morphologies or numbers over 8 days (4 IDCs) (**Figure 4B-C**). These results suggest that PfsPPase2 plays some essential functions that are not fulfilled by PfsPPase1 alone.

**Figure 4.**
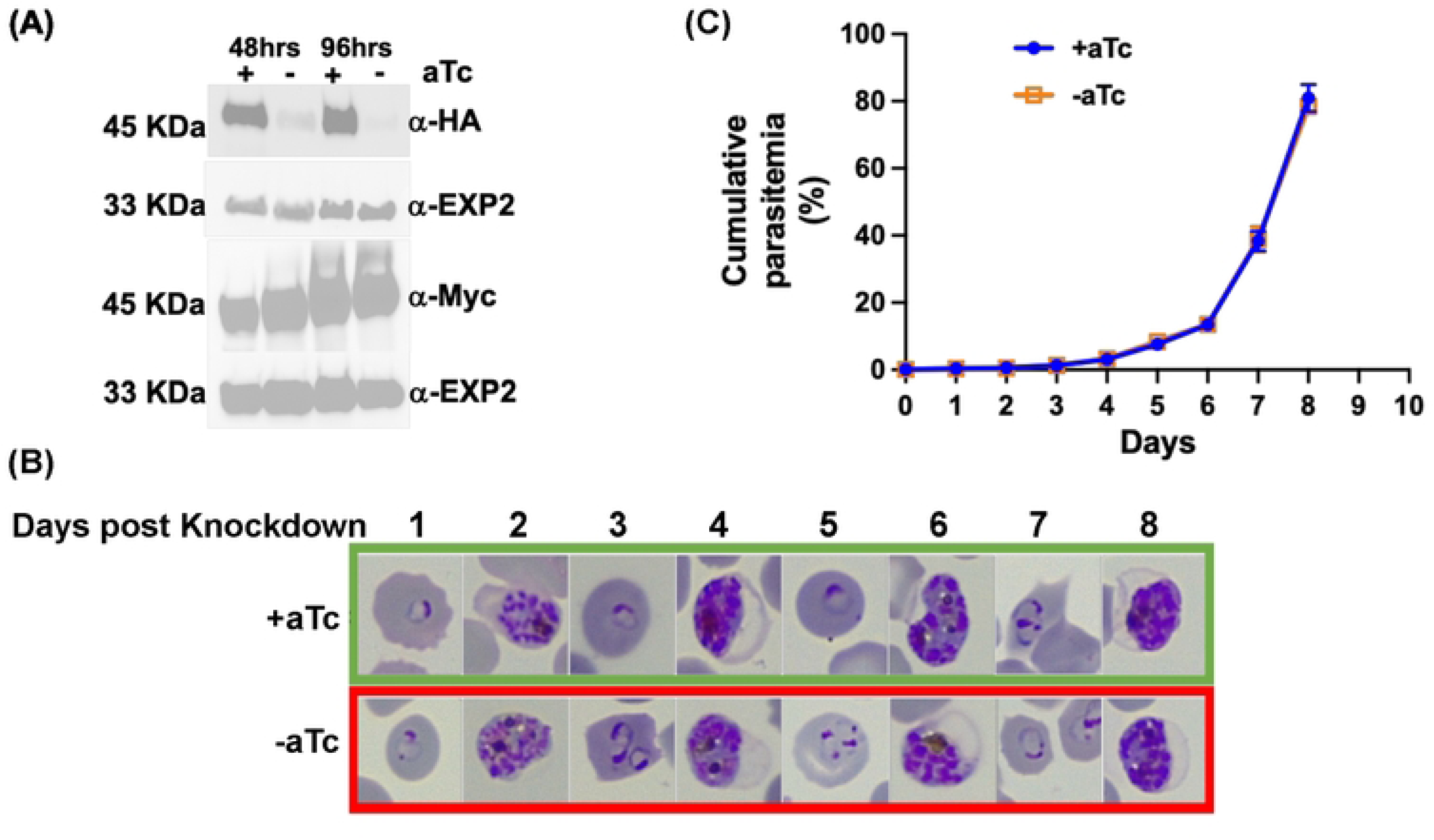
PfsPPase2 fully complements PfsPPases knockdown parasites. A, Western blot analysis of protein lysate from NF54attB-PfsPPase-3HA^APT^-PfsPPase2-3Myc. The blot was probed with anti-HA to verify knockdown of the endogenously tagged PfsPPases-3HA and re-probed with anti-PfEXP2 to show loading controls. Duplicate samples were also blotted with anti-Myc to confirm the expression of the episomally tagged PfsPPase2-3Myc and re-probed with anti-PfEXP2 to show loading controls. B, Giemsa-stained thin blood smears showing parasite morphological changes after aTc removal. Green box, aTc (+). Red box, aTc (-). C, Parasitemia of the knockdown experiment after the removal of aTc. Parasitemia was determined by microscopic counting in triplicate samples. These experiments were repeated four times.

To further verify whether PfsPPase1-3Myc and PfsPPase2-3Myc worked enzymatically properly in degrading PPi, we quantified PPi levels in the parasite lines, PfsPPase1-3Myc and PfsPPase2-3Myc, grown in aTc (±) conditions at different timepoints upon aTc removal. As shown in **Figure 2F**, the parental line NF54attB-PfsPPases-3HA^APT^ had elevated PPi levels at 96 h post aTc removal. However, at the same timepoint, the elevated PPi levels in the aTc (-) conditions were completely abolished when either PfsPPase1-3Myc or PfsPPase2-3Myc was expressed (**Figure 5**). This indicates that the episomal PfsPPase1-3Myc or PfsPPase2-3Myc was sufficient to prevent PPi accumulation when the endogenous PfsPPases were ablated. We extended our PPi measurements to 192 h (4 IDCs) of aTc removal and still observed no significant differences between the aTc (±) groups (**Supplementary Figure 4**). These results indicate that both PfsPPase1 and PfsPPase2 are enzymatically functioning in the parasites to prevent PPi accumulation.

**Figure 5.**
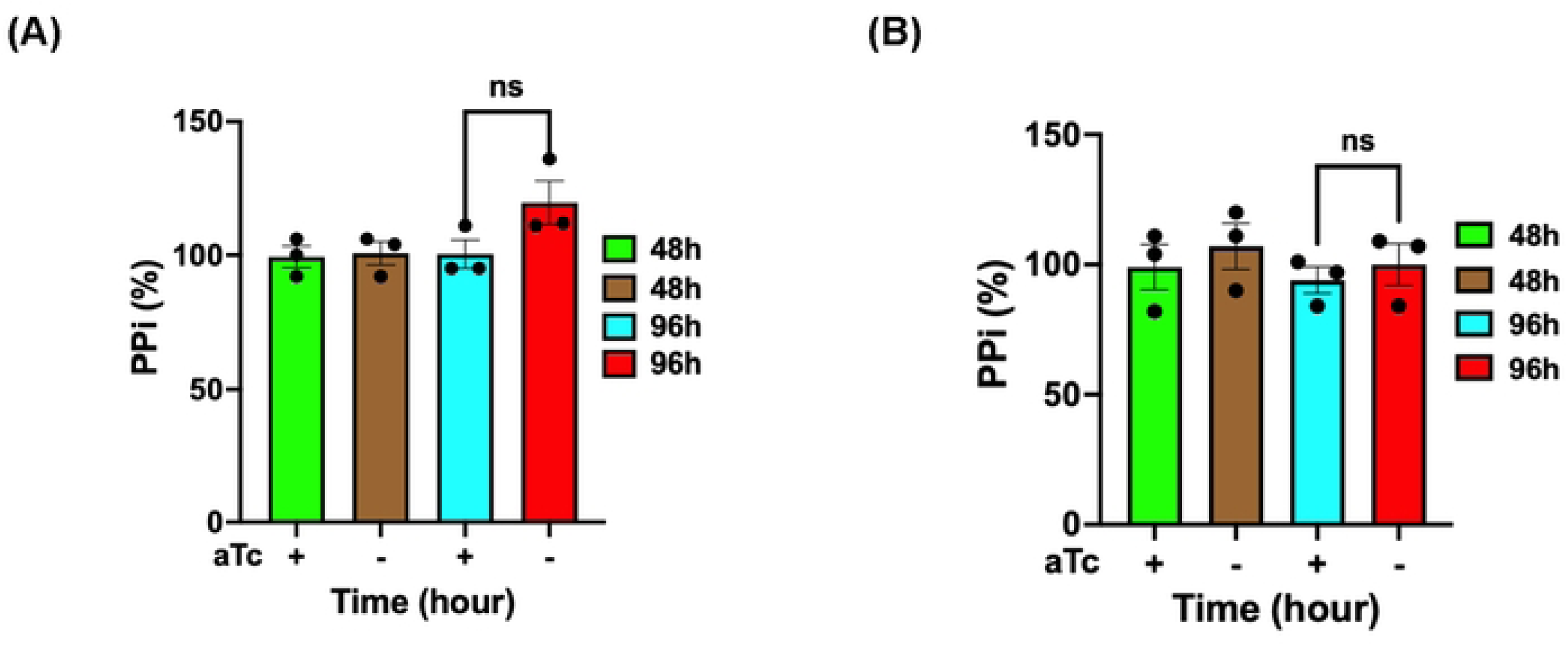
PPi levels in PfsPPase1 and PfsPPase2 complemented lines. Measurement of PPi in the PfsPPase1-3myc and PfsPPase2-3myc parasite lines after aTc removal for 48 h and 96 h from the schizont stage. PPi concentration was measured by a Sigma kit and quantified in nanomoles of PPi/mg of protein, as described previously [10]. The relative PPi concentration at various timepoints was normalized to that of the control (aTc+). A, PfsPPase1-3myc. B, PfsPPase2-3myc. Mean ± SEM of the 3 measurements is shown and statistical analysis was done by t-test. not significant (ns), p value (0.1186 and 0.5628 respectively). These experiments were repeated three times.

### Subcellular localization of PfsPPases by fixed-or live-cell fluorescence microscopy

Our results suggested that while PfsPPase1 partially rescues the loss of the endogenous PfsPPases, PfsPPase2 fully rescues. To verify their subcellular localization, we utilized both fixed-and live-cell fluorescence microscopy. In the NF54attB-PfsPPases-3HA^APT^ parasite line, we conducted immunofluorescence assay (IFA) using antibodies against HA, PfHSP60 (heat shock protein 60, a mitochondrial marker [41, 42]), and PfACP (acyl carrier protein, an apicoplast marker [30, 43]). Our data showed that PfsPPases exhibit strong diffused staining throughout the parasite, indicating that PfsPPases are predominantly localized to the parasite cytosol, in agreement with the previous study [18]. However, IFA also revealed that PfsPPases display some punctate staining patterns which partially co-localize with the structures detected by anti-PfHSP60 (**Figure 6A**) or anti-PfACP antibodies (**Figure 6B**). This result suggests that endogenous PfsPPases mainly localize to the parasite cytoplasm and additionally to the organelles (the mitochondrion and the apicoplast).

**Figure 6.**
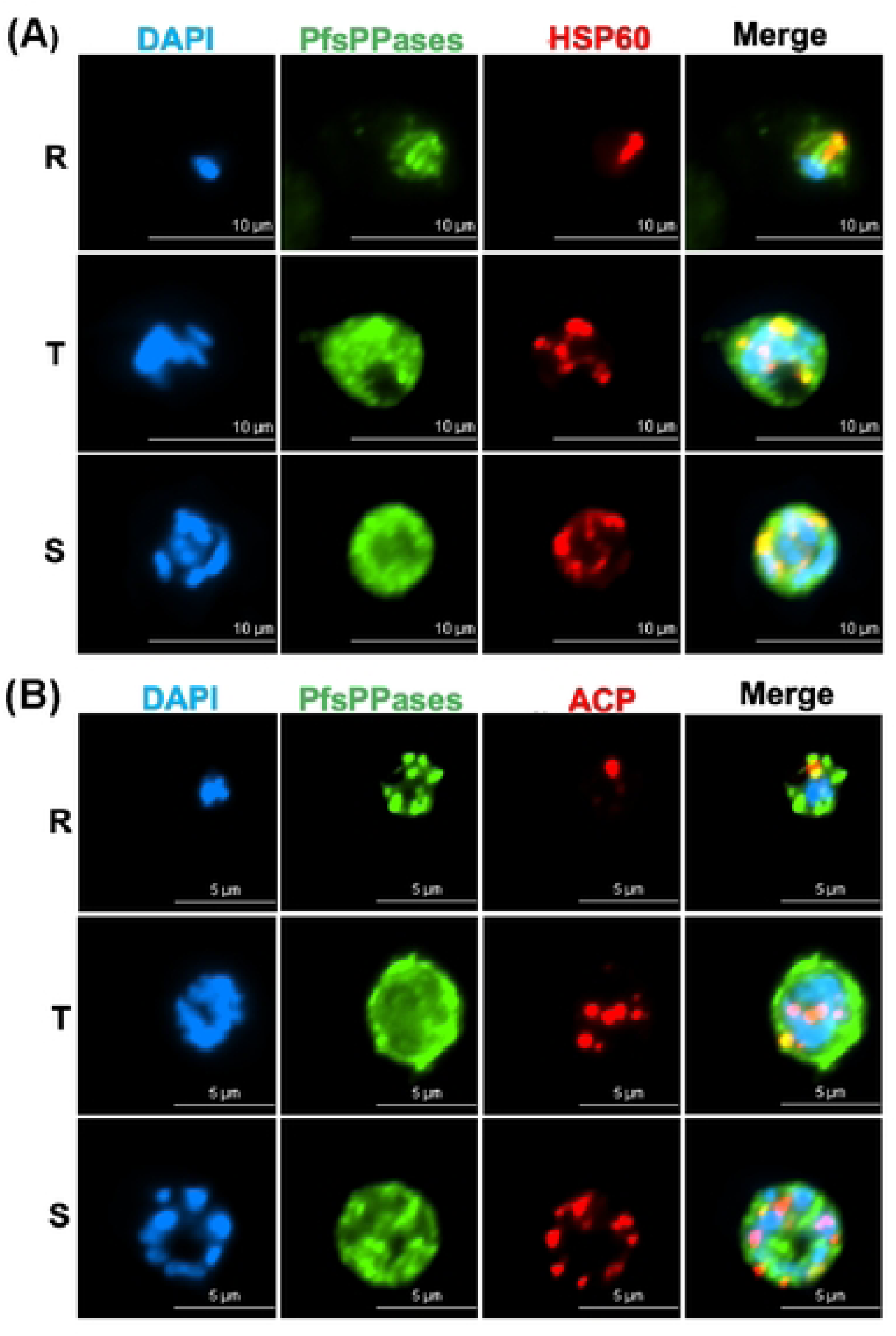
Subcellular localization of PfsPPases. A-B, Immunofluorescence assay demonstrating the expression and localization of PfsPPases-3HA throughout the parasite’s IDC. R-Ring, T-Trophozoite, S- Schizont. DAPI stained the nuclei (blue) and PfsPPases-3HA is shown in Green. A, Colocalization of PfsPPases with the mitochondrion detected by anti-PfHSP60 (red). Scale bar,10 µm. The Pearson correlation coefficient between green and red fluorescence was calculated from n = 10 parasites at each stage. Ring (0.7713 ± 0.0548), Trophozoite (0.7197 ± 0.0845), and Schizont (0.7883 ± 0.0374). B, Colocalization of PfsPPases with the apicoplast detected by anti-PfACP (red). Scale bar, 5 µm. The Pearson correlation coefficient between green and red fluorescence was calculated from n = 10 parasites at each stage. Ring (0.6493 ± 0.0632), Trophozoite (0.5979 ± 0.0781), and Schizont (0.5967 ± 0.0943).

To further explore the localization of individual isoforms, we performed IFA in the parasite lines, PfsPPase1-3Myc or PfsPPase2-3Myc, that expressed only one isoform after the endogenous 3HA-tagged PfsPPases were depleted by aTc removal for 96 h. IFA revealed that PfsPPase1 was mainly localized to the parasite cytosol with no apparent colocalization with markers against the mitochondrion or the apicoplast (**Figure 7A-B**). By contrast, PfsPPase2 exhibited strong colocalization with the mitochondrion and partial colocalization with the apicoplast (**Figure 7C-D**). Unexpectedly, PfsPPase2 was not merely organellar; it was also detected in the parasite cytosol as IFA showed some diffused staining besides the punctate structures. These results indicate that while PfsPPase1 is solely cytoplasmic, PfsPPase2 localizes to multiple compartments, including the mitochondrion, the apicoplast, and the cytoplasm. While the mitochondrial localization of PfsPPase2 corroborates well with the Mitoprot prediction (a score of 0.8), the mechanisms by which PfsPPase2 localizes to the apicoplast and cytoplasm remain unknown (see Discussion). Nevertheless, the localization patterns of PfsPPases are consistent with their functions: PfsPPase2 is fully sufficient to maintain PPi homeostasis of the entire parasite whereas PfsPPase1 is mainly responsible for the cytoplasm.

**Figure 7.**
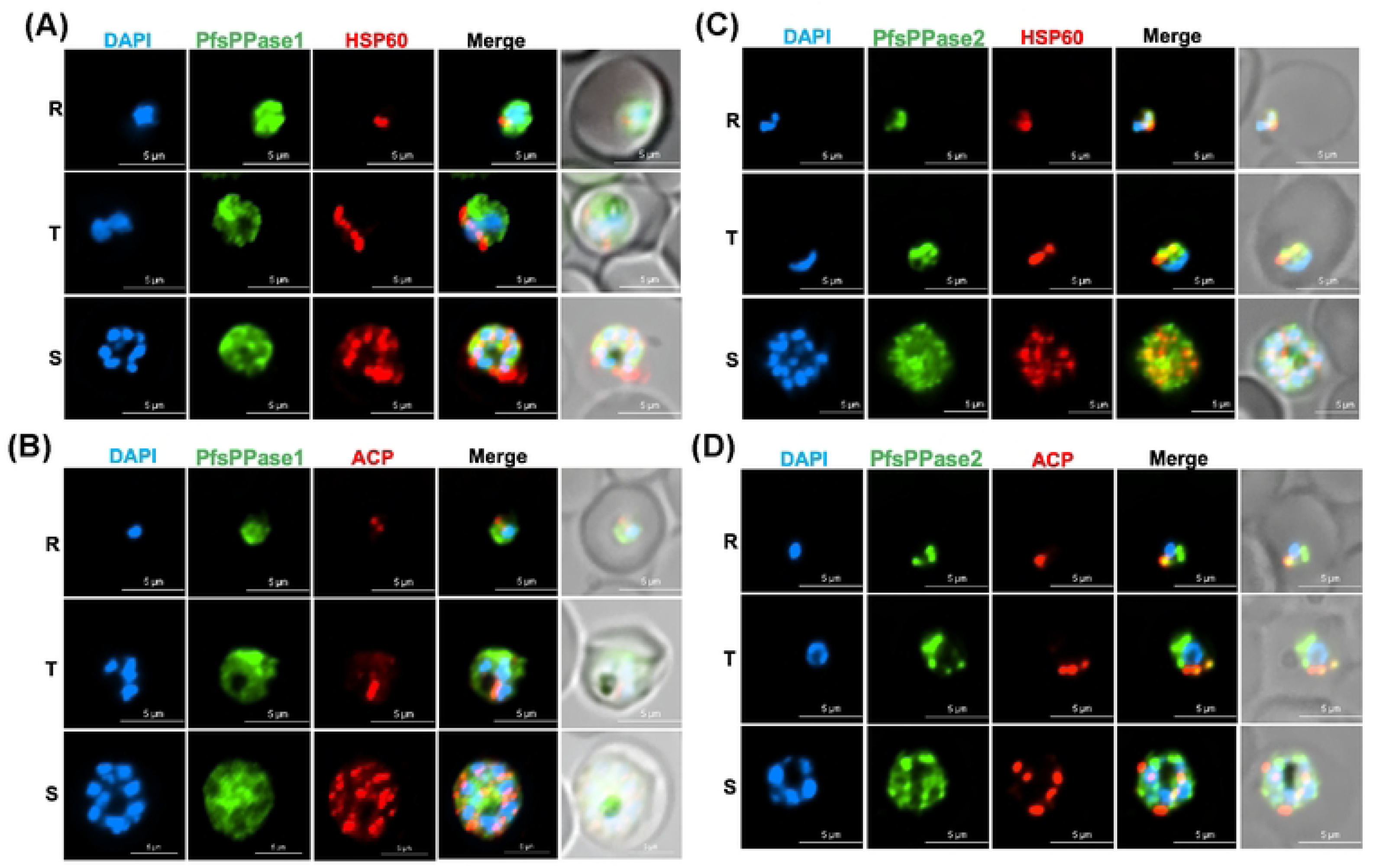
Subcellular localization of PfsPPase1 and PfsPPase2. Immunofluorescence assays of NF54attB-PfsPPase-3HA^APT^-PfsPPase1-3Myc (A-B) or NF54attB-PfsPPase-3HA^APT^-PfsPPase2-3Myc (C-D), conducted at 96 h post aTc removal to deplete the endogenous PfsPPases. The samples were verified by anti-Myc and anti-PfHsp60 (A, C) or by anti-Myc and anti-PfACP (B, D). PfHSp60 is a mitochondrial marker; PfACP is an apicoplast marker. Scale bars, 5 μm. In A-D, Pearson correlation coefficient of green and red fluorescence was derived from n = 10 parasites of each stage. The values are as follows, A, Ring (0.4791 ± 0.1082), Trophozoite (0.4591 ± 0.0873), and Schizont (0.4656 ± 0.0744); B, Ring (0.3977 ± 0.0632), Trophozoite (0.4132 ± 0.0612), and Schizont (0.4483 ± 0.0552); C, Ring (0.7729 ± 0.0429), Trophozoite (0.7532 ± 0.0537), and Schizont (0.7475 ± 0.0643); and D, Ring (0.6834 ± 0.0496), Trophozoite (0.6753 ± 0.0590), and Schizont (0.6190 ± 0.0602).

To validate the IFA results with fixed parasites, we generated two new parasite lines that express endogenous PfsPPases tagged with mRuby for live-cell microscopy. The parasite lines were named NF54attB-PfsPPases-mRuby^APT^ and PfMev-PfsPPases-mRuby^APT^ with the latter contains a GFP labeled apicoplast [44]. Using the NF54attB-PfsPPases-mRuby^APT^ parasite line, we stained the parasite’s mitochondrion with MitoTracker and observed strong fluorescence of PfsPPases in the mitochondrion and cytosol across all stages of the parasite’s IDC, confirming the localization of PfsPPases to both the cytosol and mitochondrion (**Figure 8A**). In the PfMev-PfsPPases-mRuby^APT^ parasite line, live-cell microscopy detected fluorescence of PfsPPases in the cytoplasm and, to a varying extent, in the apicoplast across different stages of the parasite’s IDC (**Figure 8B**). Further, quantification of PfsPPases localization revealed that nearly 100 percent of the ring stage parasites exhibited PfsPPases in the apicoplast, while 60 percent of trophozoite stage parasites, and 80 percent of schizont stage parasites showed some degrees of PfsPPases localization in the apicoplast. Collectively, our live-cell microscopy data reveal that PfsPPases are localized to the cytosol, the mitochondrion and the apicoplast.

**Figure 8.**
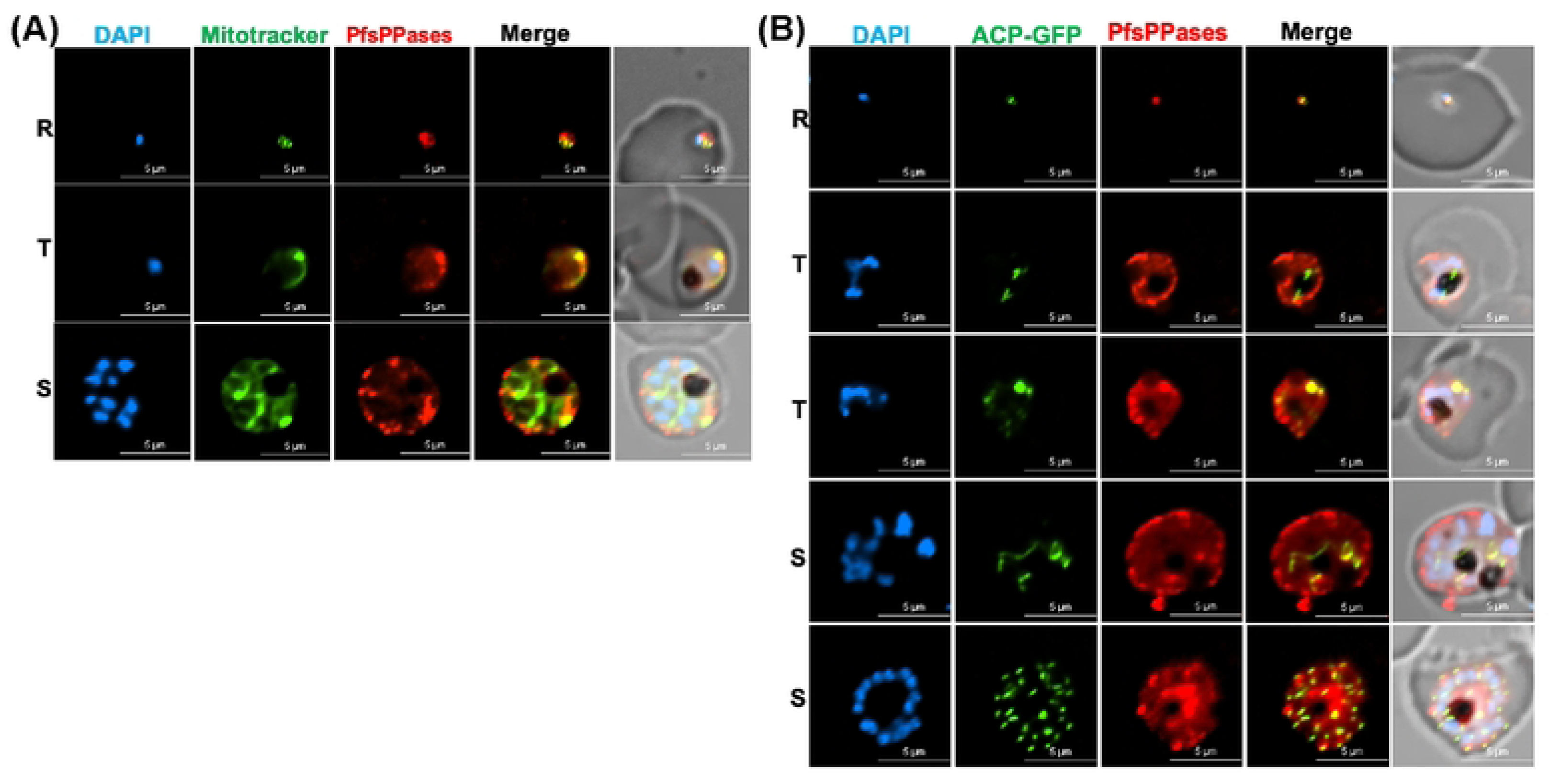
Subcellular localization of PfsPPases via live microscopy. Live imaging of NF54attB-PfsPPases-mRuby showing expression and localization of the enzymes throughout the parasite’s IDC. R-Ring, T-Trophozoite, S-Schizont. DAPI stained the nuclei. Red, PfsPPases-mRuby. A, Colocalization of PfsPPases with the mitochondrion detected by green MitoTracker. Scale bars, 10 μm. Pearson correlation coefficient of green and red fluorescence was derived from n = 10 parasites of each stage. Ring (0.8967 ± 0.0489), Trophozoite (0.8252 ± 0.0432), and Schizont (0.8303 ± 0.0563). B, Colocalization of PfsPPases with the apicoplast labeled by GFP (guided by the first 55-aa of acyl carrier protein). Scale bars, 5 μm. Pearson correlation coefficient of green and red fluorescence was derived from n = 10 parasites of each stage. Ring (0.8609 ± 0.0664), Trophozoite (0.7657 ± 0.0833), and Schizont (0.8000 ± 0.0766).

To further verify the organellar localization of PfsPPase2, we generated another parasite line (NF54attB-PfsPPase2^leader^-mNeonGreen) in which the 51-aa leader sequence of PfsPPase2 was fused with mNeonGreen (mNG) and the fusion gene was integrated into a non-essential genomic locus, cg6 [45], and expressed under the mRL2 promoter. Using the NF54attB-PfsPPase2^leader^-mNeonGreen parasite line in live-cell microscopy, we stained it with MitoTracker and confirmed that the leader sequence of PfsPPase2 is sufficient to direct the protein to the mitochondrion (**Supplementary Figure 5A**). Again, this result corroborates well with the high predicted score of PfsPPase2 for mitochondrial localization (0.8 by MitoProt). Moreover, we also investigated the potential apicoplast targeting of the PfsPPase2 leader sequence by IFA using anti-PfACP. IFA suggested that PfsPPase2 is localized to the apicoplast in 50% of parasites that were examined (**Supplementary Figure 5B**). Thus, it appears that the 51-aa leader sequence is sufficient to guide PfsPPase2 to the organelles, including the mitochondrion and the apicoplast.

### Subcellular localization of PfsPPases by a cytoplasmic protein leakage assay

As shown above, we have utilized both fixed-and live-cell microscopy to verify the subcellular localization of PfsPPases (**Figures 6-8 and Supplementary Figure 5**), revealing that while PfsPPase1 is cytoplasmic, PfsPPase2 can localize to the mitochondrion, apicoplast, and cytoplasm. To further verify this complex localization pattern of PfsPPases, we took advantage of a cytoplasmic protein leakage assay that was previously developed by the Vaidya group. As shown by Das *et al.* [46], cytoplasmic proteins, but not organellar or membrane-associated ones, leak out from the parasites upon treatments of inhibitors against PfATP4, a sodium pump, and saponin. Once PfATP4 is inhibited, saponin becomes effective on the plasma membrane parasite (PPM) of the treated parasites as cholesterol likely accumulates on their PPM [46]. Upon leakage, cytoplasmic proteins such as aldolase are no longer detectable in the parasite pellet. We reasoned that this cytoplasmic protein leakage assay is an ideal complementary approach to our microscopic data to verify the subcellular localization of PfsPPases.

We first treated the NF54attB-PfsPPases-3HA^APT^ line with KAE609 (a PfATP4 inhibitor [47, 48]) for 2 h then performed saponin treatment. The samples were then analyzed by western blot to detect PfsPPases-3HA in the parasite pellet. Two control proteins were used to validate the assay, including the cytosolic PfAldolase (should be leaked) and the membrane-bound PfExp2 (should be retained). Western blot revealed that KAE609 treatment renders a complete leakage of PfAldolase, but no loss of PfExp2, ensuring that the protein leakage assay worked properly (**Figure 9A-B; Supplementary Figure 6A-C**). PfsPPases were observed to lose partially in the parasite pellet in a concentration dependent manner, but some protein remained even at the highest KAE609 concentration, suggesting that PfsPPases are cytoplasmic as well as organellar. To further validate the distinct localization of each PfsPPase isoform, we repeated the experiment using PfsPPase1-3Myc and PfsPPase2-3Myc lines at 96 h post aTc removal to deplete the 3HA-tagged endogenous PfsPPases. Post KAE609 treatment (2 h) and saponin lysis, the pellet samples were analyzed by western blot, which revealed that PfsPPase1 was partially lost at a low concentration of KAE609 but was totally leaked out at the highest concentration. This strongly indicated that PfsPPase1 is exclusively cytosolic (**Figure 9C-D; Supplementary Figure 6D-F)**. In contrast, PsPPase2 showed no significant loss at a low concentration of KAE609 treatment and only a minor loss at the highest concentration (**Figure 9C, 9E; Supplementary Figure 6D-F**), indicating that PfsPPase2 localizes to organelles as well as cytoplasm. Collectively, in combination with our fixed-and live-cell microscopic data, this cytoplasmic protein leakage assay strongly suggests that while PfsPPase1 is exclusively cytosolic, PfsPPase2 has cytosolic and organellar localization, further supporting the functional differences of the two PfsPPase isoforms.

**Figure 9.**
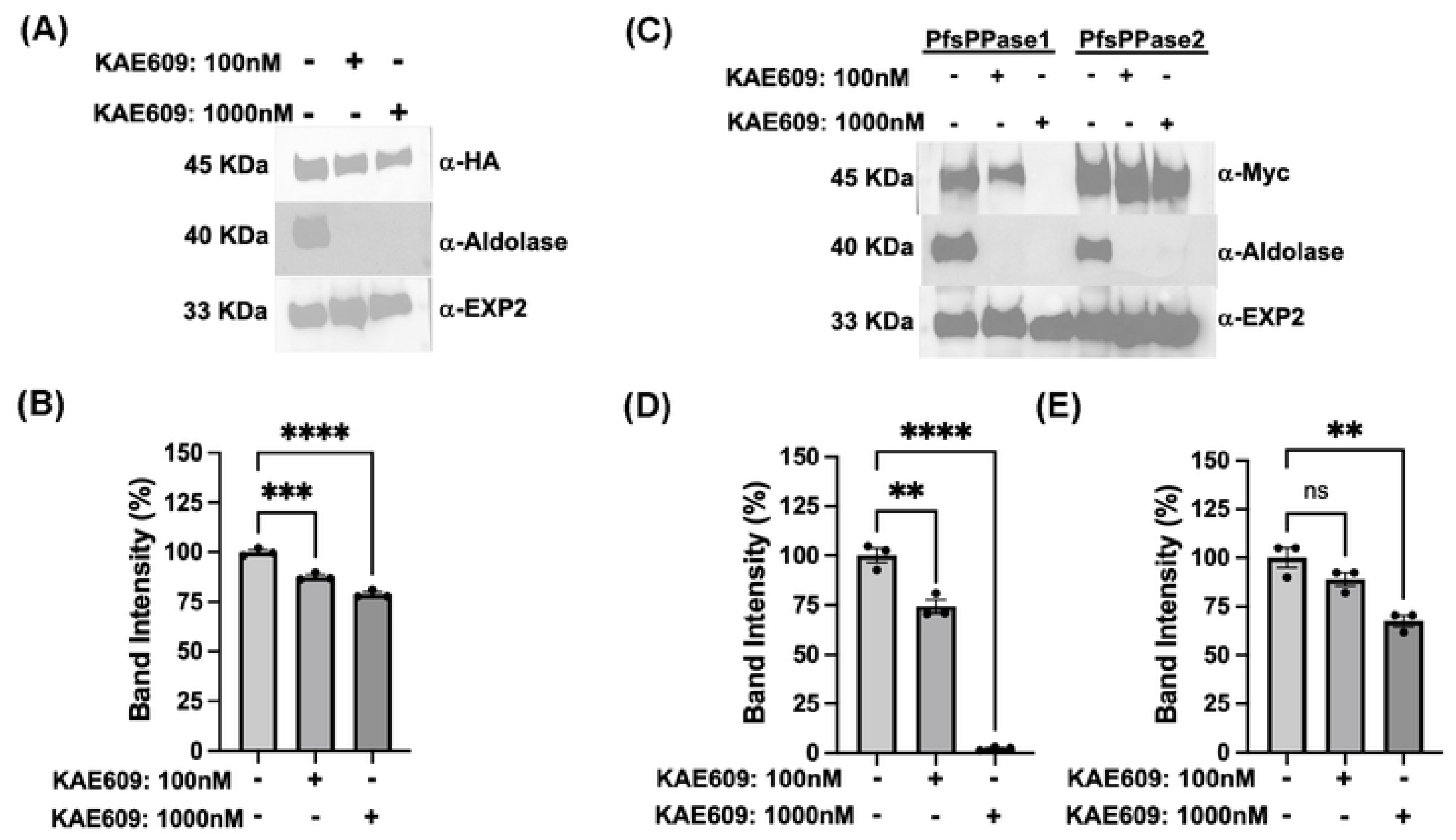
A cytoplasmic protein leakage assay shows PfsPPase1 is solely cytosolic while PfsPPase2 localizes to organelles and cytoplasm. A, Western blot analysis of NF54attB-sPPase-3HA^APT^ treated with 0, 100 or 1000 nM KAE609 for 2 h. The samples were probed with anti-HA, anti-PfAldolase, or anti-PfEXP2. B, Band intensity measurement of A and two additional replicates with Image J. C-D, Western blot analysis of PfsPPase1-3Myc and PfsPPase2-3Myc parasites treated with 0, 100 or 1000 nM KAE609 for 2 h. D, Band intensity measurement of PfsPPase1-3Myc using image J. E, Band intensity measurement of PfsPPase2-3Myc using image J. Statistical analysis was done by t-tests. **, *p* < 0.05; ***, *p <* 0.001, ****, *p <* 0.0001; ns, non-significant.

## DISCUSSION

In this study, we employed reverse genetics and biochemical approaches to characterize soluble pyrophosphatases in *Plasmodium falciparum*, elucidating their functional roles during asexual blood stages. Our data indicate that PfsPPases are essential for parasite growth and development. Despite being expressed throughout the asexual blood stages [18], knockdown of PfsPPases results in parasite arrest at the trophozoite stage, preventing its progression to the schizont stage (**Figure 2**). We have previously demonstrated that the proton-pumping vacuolar pyrophosphatase (PfVP1) is essential in the ring stage [10]. Together, our findings highlight that PPi homeostasis is essential in all asexual blood stages, with PPi hydrolysis mediated by distinct mechanisms—either via membrane-bound or soluble pyrophosphatases—at various stages of the 48-h IDC. Interestingly, despite being expressed in the late stages, PfVP1 is unable to compensate for the loss of PfsPPases. This suggests that the increased metabolic activities of the trophozoite stage [49–51] necessitates robust PPi degradation mediated by soluble pyrophosphatases. Although the previous study characterized the enzymatic activities of PfsPPases in *P. falciparum* [18], our study is the first to report the essentiality of these enzymes and characterize the differences of two PfsPPase isoforms.

We have utilized multiple approaches to localize PfsPPases to the parasite cytoplasm, the mitochondrion and potentially the apicoplast (**Figures 6-8, Supplementary Figure 5**). This localization pattern aligns with the presence of three distinct genomes located in the nucleus (23 Mb) [22, 23], the mitochondrion (6 kb) [24–26], and the apicoplast (35 kb) [27–30], suggesting that PPi homeostasis is critical in all these subcellular compartments (see a model in **Figure 10**). Although replication of the nuclear genome and translation of several thousand proteins would generate high levels of PPi, PPi is also generated during DNA replication, gene transcription, protein synthesis and other cellular processes occurring within the mitochondrion and the apicoplast. Nuclearly generated PPi can be trafficked/diffused through the nuclear pores into the cytoplasm, where it is degraded by cytosolic pyrophosphatase alongside PPi generated from cytoplasmic processes, highlighting the importance of cytoplasmic pyrophosphatases. On the other hand, PPi generated within the mitochondrion and the apicoplast needs to be hydrolyzed locally to prevent its toxic accumulation as no PPi transporters are annotated in the parasite genome, which aligns with the fact that PPi transporters have not been reported in model organisms [7, 16]. Our data underscore the critical role of two PfsPPases in maintaining pyrophosphate homeostasis in the parasite’s cytosol and organelles (mitochondrion and apicoplast).

**Figure 10.**
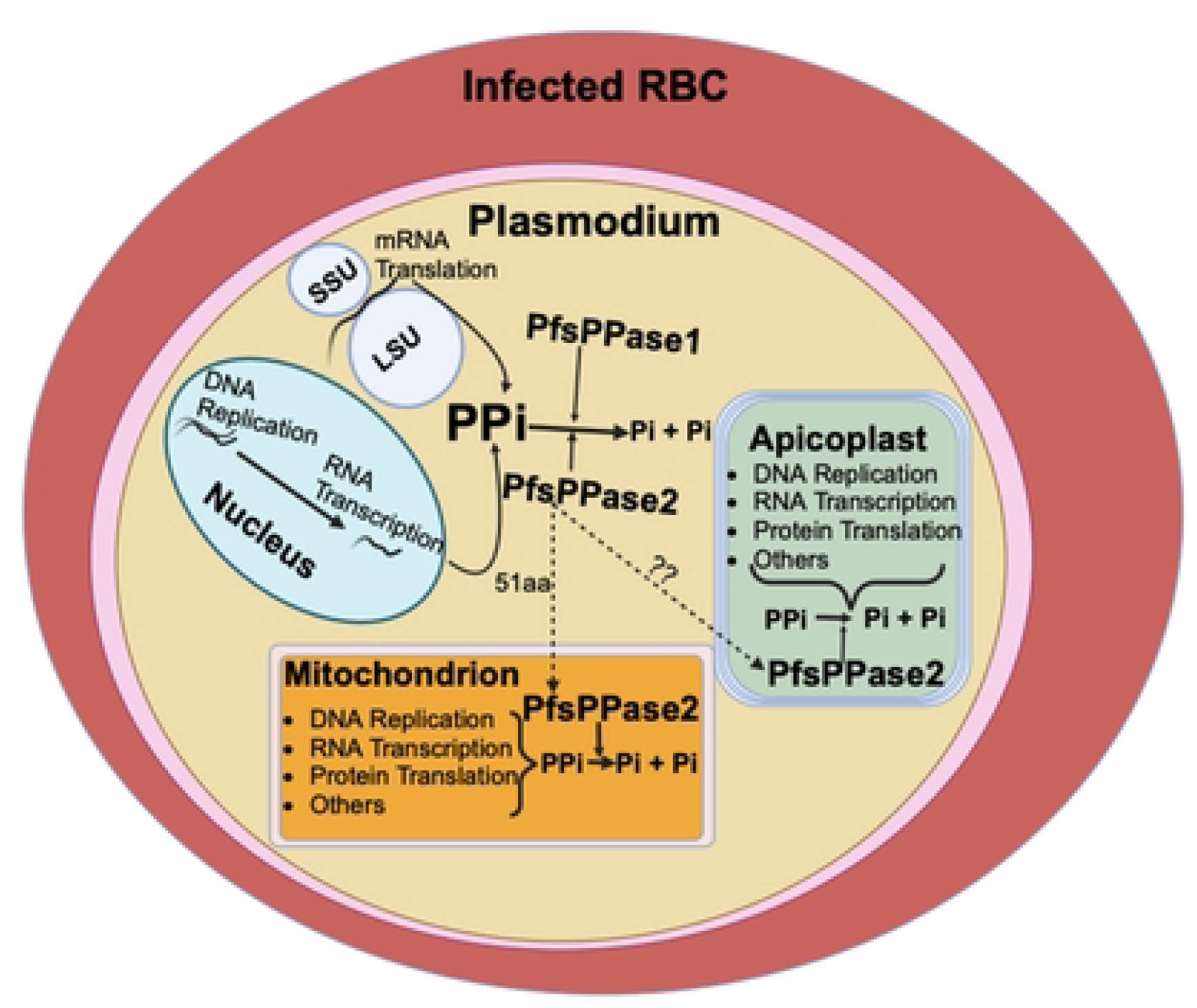
**Schematic representation depicting the localization and functions of two PfsPPase isoforms in *Plasmodium falciparum***. PfsPPase1 is localized solely to the parasite cytoplasm while PfsPPase2 displays a complex localization pattern, being present in the organelles (the mitochondrion and the apicoplast) and the cytosol. Two isoforms of PfsPPases work together to maintain pyrophosphate homeostasis in multiple subcellular compartments of *P. falciparum*. Question marks represent unknown mechanisms.

To the best of our knowledge, this is the first study to characterize two isoforms, PfsPPase1 and PfsPPase2, encoded by the same gene locus (Pf3D7_0316300) (**Figure 1**). The key difference between these isoforms is the presence of an additional 51-aa leader sequence in PfsPPase2 (**Figure 1A**). RNA-seq clearly indicates that the two short exons upstream in the gene locus are transcribed, encoding most of the 51-aa leader peptide (**Supplementary Figure 1**). The leader sequence is predicted to direct PfsPPase2 to the mitochondrion (MitoProt score of 0.8), which was further validated in this study (**Supplementary Figure 5**). Although PlasmoAP, an apicoplast prediction tool, does not predict PfsPPase2’s localization in the apicoplast, we also showed that PfsPPase2 partially localizes to the apicoplast (**Figures 6-8, Supplementary Figure 5**). Interestingly, both isoforms are independently transcribed but are not the products of alternative splicing from a single gene (**Figure 1**). Further, western blot analysis of PfsPPases only detects one protein band at the molecular weight of PfsPPase1, suggesting that a rapid cleavage of the 51-aa leader sequence from PfsPPase2 likely occurs during protein translocation, although the exact cleavage site and mechanism remain unknown. Notably, the previous study focused on the structural characterization of sPPases in Apicomplexa but did not recognize the existence of two PfsPPase isoforms in *P. falciparum* [18]. Hence, the organellar localization of PfsPPases was likely overlooked in IFA studies that utilized homemade polyclonal antibodies in wildtype parasites [18]. By contrast, in this study, we employed endogenous tagging and multiple approaches to confirm the localization of both PfsPPase isoforms in six transgenic parasite lines with either the full-length protein or the 51-aa leader sequence tagged by various epitopes, including 3HA, mRuby, or mNeonGreen. Our results showed that while PfsPPase1 is solely cytoplasmic, PfsPPase2 exhibits a complex localization pattern, being present in the mitochondrion, the apicoplast, and the cytoplasm. Moreover, the microscopic localization data are well supported by the results of the cytoplasmic protein leakage assay shown in **Figure 9**. Altogether, using multiple approaches, we conclude that the gene locus of PfsPPase (Pf3D7_0316300) encodes two isoforms that maintain PPi homeostasis in the parasite cytoplasm, the mitochondrion and the apicoplast. Notably, the mechanisms by which the same genetic locus encodes two PfsPPase isoforms remain unclear and deserve to be further investigated.

Using a combination of genetic knockdown and complementation studies, for the first time, we have elucidated the functional differences of two PfsPPase isoforms. PfsPPase1 is mainly localized to the parasite cytoplasm (**Figures 6-8**) and complementation of the PfsPPases knockdown parasites with PfsPPase1 alone did not fully rescue parasite growth, with abnormal morphologies observed by day 6 post aTc removal (**Figure 3**). By contrast, PfsPPase2 is localized to the mitochondrion, the apicoplast, and the parasite cytoplasm (**Figures 6-8, Supplementary Figure 5**). Furthermore, complementation of the PfsPPases knockdown parasites with PfsPPase2 alone fully rescued their growth with no phenotypic defects observed (**Figure 4**).

These results imply that organellar PPi hydrolysis, in addition to cytoplasmic degradation of PPi, is critical for maintaining the parasite intracellular equilibria. Therefore, we speculate that the reasons why PfsPPase1 alone was unable to fully rescue the loss of PfsPPases are due to (i) the absence of the 51-aa leader sequence in PfsPPase1, (ii) its confinement to the parasite cytoplasm alone, and (iii) the resultant accumulation of PPi in the organelles, which could compromise their integrity and ultimately lead to parasite death. Notably, when we quantified PPi levels in parasites complemented with either PfsPPase1 or PfsPPase2 at 96 h or 192 h post aTc removal, we did not observe any significant PPi accumulation in the knockdown parasites (**Figure 5, Supplementary Figure 4**). While these results suggest that PfsPPase1 or PfsPPase2 is enzymatically functional in the parasites, it does not directly show organellar accumulation of PPi in the PfsPPase1 complemented parasites. It is important to note that the majority of PPi generated within the parasites should be present in the cytoplasm and the organellar PPi levels remain unspecified in our protocol due to the lack of established methods to completely purify mitochondria and apicoplasts from *P. falciparum*. Clearly, future studies are needed to confirm the accumulation of PPi in these organelles in the absence of organellar PfsPPases.

While PfsPPase2 is bioinformatically predicted to be mitochondrial, it is interesting to note that PfsPPase2 also localizes to the apicoplast without any bioinformatic evidence for the presence of an apicoplast localization signal or peptide. This raises an intriguing question of how the enzyme is targeted to the apicoplast. The answers to this remain unclear at present. We speculate that PfsPPase2 is trafficked to the apicoplast either directly through the 51-aa leader sequence as shown in **Supplementary Figure 5**, via interactions between the mitochondrion and the apicoplast contact sites, or by an unknown mechanism(s). Recently, it has been shown in *Toxoplasma gondii*, an apicomplexan parasite distantly related to *P. falciparum*, that proteins with dual localization in both the apicoplast and the mitochondrion are translocated via a Golgi-dependent pathways [52]. Whether or not similar mechanisms apply to PfsPPase2 deserves to be further investigated. Unexpectedly, our results also show that PfsPPase2 is partially cytoplasmic (**Figures 7,9**). The protein leakage assay suggests that a portion of PfsPPase2 is cytoplasmic and leaks out upon treatments of PfATP4 inhibitors and saponin, whereas most of the enzyme is associated with organelles (**Figure 9**). Despite this, the full-length PfsPPase2 at 51 kDa was undetectable, suggesting that cytoplasmic PfsPPase2 is either below the detection limit or is also cleaved to remove the 51-aa leader sequence. Future studies are needed to elucidate the mechanisms of PfsPPase2 protein trafficking within the parasite.

Our discovery of cytoplasmic and organellar localization of PfsPPases is consistent with the significance of soluble pyrophosphatases in other organisms. It has been long recognized that *S. cerevisiae* and mammals have two soluble pyrophosphatases located in the cytoplasm and the mitochondrion [15, 16, 19]. Interestingly, these two soluble pyrophosphatases are encoded by different genes and only share moderate sequence identity (47% and 62% identify for yeast [19] and human [15] enzymes, respectively). In *Arabidopsis thaliana*, the cytoplasm contains five soluble pyrophosphatase isoforms with 69% identity (one of which is also mitochondrial) and one plastid pyrophosphatase that is highly divergent from all other counterparts [20, 21]. In comparison to these model organisms that encode distinct genes to ensure sufficient soluble pyrophosphatases in various compartments, *P. falciparum* has seemingly taken a distinct route to duplicate the enzymes by adding a leader sequence in front of the soluble pyrophosphatase. The presence of the additional 51-aa leader sequence ensures the targeting of PfsPPase2 to the organelles (mitochondrion and partially, apicoplast) for degradation of locally generated PPi. It may appear that the presence of two isoforms of sPPases from the same gene locus is unique to *P. falciparum* as this genetic configuration was not evidently observed in other apicomplexan parasites such as *Toxoplasma gondii* [53]. Since *T. gondii* contains essential functions in both the mitochondrion and the apicoplast, it would be unlikely that PPi homeostasis in these organelles is not essential. Therefore, it is plausible that *T. gondii* and other apicomplexan parasites also translocate sPPases into organelles, although the exact mechanisms remain entirely unknown.

In summary, our data suggest that degradation of PPi generated in both the cytoplasm and organelles (the mitochondrion and the apicoplast) is essential for maintaining the overall cellular homeostasis of *P. falciparum*. This process is crucial for the growth and development of the parasite during the late phases of the asexual blood stage, particularly at the transition from the trophozoite stage to the schizont. The essential nature of PfsPPases, coupled with their divergence from the human orthologs, makes them considerable drug targets against malaria.

## Materials and methods

### Plasmid construction

1, Endogenous tagging of PfsPPases with 3HA (hemagglutinin). To generate the transgenic parasite line NF54attB-PfsPPases-3HA^APT^, we modified the endogenous locus of PfsPPases (PF3D7_0316300) via CRISPR/Cas9 [33, 34]. To construct the pMG75 vector plasmid, we amplified two homologous regions (5HR and 3HR) from *P. falciparum* genomic DNA using primers P7-P10 (Supplementary Table 1). These two PCR products were annealed using an overlap extension PCR [54], forming a 3HR+5HR fused fragment. It was then digested with AflII and BstEII, cloned into the pMG75-3HA plasmid [55], and sequenced with the vector primers (P11-P12), yielding the pMG75-PfsPPases-3HA plasmid. For parasite transfection, we linearized the plasmid with SacII and mixed it with the circular gRNA plasmid. The gRNA sequence was selected using the Eukaryotic Pathogen CRISPR guide RNA design tool (http://grna.ctegd.uga.edu/) and cloned into our NF-Cas9 plasmid [55], which contained the Cas9 gene from *Streptococcus pyogenes*. gRNA cloning was done via the NEB HiFi-DNA Assembly Master mix as described previously [55], and the gRNA sequence was confirmed by Sanger sequencing using primer (P14).

2, Endogenous tagging of PfsPPases with mRuby. We used the previously PCR-amplified PfsPPases 3HR+5HR fragment, digested it with AflII and BstEII, and cloned it into the pMG75-PfVP1-mRUBY plasmid (a derivative of pMG75-PfVP1-mNeonGreen, [10]). The correct insertion of 3HR+5HR was confirmed by sequencing using primer P15.

3, Generation of NF54attB-PfsPPases-3HA^APT^-PfsPPase1-3Myc and NF54attB-PfsPPases-3HA^APT^-PfsPPase2-3Myc parasite lines. In the NF54attB-PfsPPases-3HA^APT^ line, we performed a second transfection to complement the knockdown of PfsPPases with either PfsPPase1-3Myc or PfsPPase2-3Myc, driven by the mRL2 promoter (PF3D7_1132700) [40]. Briefly, we amplified the full-length coding regions of PfsPPase1 or PfsPPase2 from parasite cDNA using primers P4 & P6 or P5 & P6, and then individually cloned them into the PLN-RL2-hDHFR-3Myc construct [39] via AvrII and BsiWI. The resulting plasmids, PLN-RL2-hDHFR-PfsPPase1-3Myc and PLN-RL2-hDHFR-PfsPPase2-3Myc, were sequenced using the PLN vector primers (P16-P17).

4, Cloning the 51-aa leader sequence for localization studies. The fragment encoding the 51-aa leader sequence of PfsPPase2 was PCR amplified from parasite cDNA using primers P20 and P21. It was then cloned into the PLN-RL2-Hsp60L-mNeongreen plasmid [56] by AvrII and NheI to replace the Hsp60 leader sequence. The correct insertion of the 51-aa leader sequence was confirmed by Sanger sequencing.

### Parasite culture, transfection, and knockdown studies

*P. falciparum* parasites were cultured in human O^+^ RBCs using RPMI-1640 media supplemented with 0.3% Albumax I as previously described [39, 55]. Ring-stage parasites (∼ 5% parasitemia) were transfected with either linearized or circular plasmids (50 µg) using a Bio-Rad electroporator. Forty-eight hours post-transfection, parasite cultures were maintained in media containing the appropriate drugs for selection, including blasticidin (2.5 µg/mL, InvivoGen) or WR99210 (5 nM, a kind gift from Jacobs Pharmaceutical). For parasites that need anhydrotetracycline (aTc) (250 nM, Fisher Scientific), aTc was added on the day of transfection. For knockdown studies, parasites were synchronized via several rounds of alanine/HEPES (0.5M/10 mM) treatment, isolated by Percoll at late stages, washed three times with 1xPBS to remove residual aTc (and Percoll), diluted with fresh RBCs, and cultured in media with or without aTc for several IDCs.

### Western blot and Protein leakage assay with PfATP4 inhibitors

Infected RBCs were treated with 0.05% Saponin/PBS supplemented with 1x protease inhibitor cocktail and protein extraction was performed using 2% SDS/62mM Tris-HCl (pH 6.8) as described previously [55]. Following high-speed centrifugation (13,000 rpm, 10 min), the supernatant was used for protein electrophoresis. After electrophoresis, the proteins were transferred onto the blot, blocked with 5% non-fat milk in PBS and incubated with primary antibodies as follows: anti-HA (1:10,000, mouse, sc-7392, Santa Cruz Biotechnology), anti-Myc (1:8,000, rabbit, 2278S, Cell Signaling), and anti-*Pf*EXP2 (1:10,000, rabbit, a gift from Dr. James Burns, Drexel University). The secondary antibodies used were HRP conjugated goat anti-mouse (A16078, ThermoFisher Scientific) at 1:10,000 and HRP conjugated goat anti-rabbit (31460, ThermoFisher Scientific) at 1:10,000. Subsequent steps followed the standard protocols. Blots were incubated with the Pierce^TM^ ECL substrates and visualized using the ChemiDoc Imaging Systems (Bio-Rad). Protein concentration for all samples was determined using the detergent-tolerant Pierce^TM^ BCA Protein Assay Kit (23227, ThermoFisher) according to the manufacturer’s instructions.

In each condition of the cytoplasmic protein leakage assay, approximately 10˄^8^ of infected RBCs were treated with various concentrations of KAE609, a PfATP4 inhibitor, for 2 h. The parasites were then treated with 0.05% Saponin/PBS supplemented with 1xprotease inhibitor cocktail, centrifuged and the pellets were used for western blot after they were lysed with 2%SDS/62 mM Tris (pH 6.8). Primary antibodies used in western blot included: anti-HA, anti-PfAldolase (cat no. ab38905, Abcam), anti-PfExp2. Other procedures followed the western blot protocols described above.

### Immunofluorescence assays (IFA)

IFA was performed as described previously [39, 55]. The following primary antibody was used: anti-HA (1:500, mouse sc-7392, Santa Cruz Biotechnology), anti-Myc (1:300, mouse 2276S, Cell Signaling), anti-PfACP (1:500, a gift from Dr. Sean Prigge, Johns Hopkins University), and anti-PfHSP60 (1:500, rabbit, NBP2-12734, Novus). Fluorescently labelled secondary antibodies for FITC or TRITC were used to incubate the samples for 3 h at room temperature. Other procedures followed the standard IFA protocol. Images were captured using the Nikon Ti microscope and processed by the NIS-Elements software.

### Live cell fluorescence microscopy

An aliquot of infected RBCs (250 µL with 5% hematocrit and 5% parasitemia) was stained with Hoechst (1:1000 dilution of 1mg/mL), added into a glass-bottomed dish (P35G-1.5-14-C, Matteck) and incubated for 30 min. The dish was pre-treated with poly-lysine (P8920, Sigma) either at 4°C for overnight or at 37°C for 1-2 h. After incubation in the dish, the extra non-adhered infected RBCs were washed off and the adhered cells were added with 1 mL of phenol-free RPMI and then imaged under the Nikon Ti microscope.

### PPi extraction and measurement

The PPi in the parasites was extracted in accordance with the published protocol with some modifications [53]. Saline/glucose buffer (NaCl 125 mM, KCl 5 mM, MgCl_2_ 1 mM, glucose 20 mM, HEPES 25mM, pH 7.4) was used for saponin lysis and washing. In each condition, 2×10˄^8^ of parasitized RBCs (or uninfected RBCs as a control) were saponin lysed and washed 3 times to remove hemoglobin followed by the addition of 2 volumes of saline/glucose buffer. The mixture was heated at 90°C for 10 min to inactivate soluble pyrophosphatases and saved at - 80°C. The samples were thawed and underwent 3 cycles of freezing/thawing between dry ice (10 min) and 37°C (∼ 2 min) and then, sonicated for 30 min at 4°C in a water bath sonicator (Fisher). After sonication, samples were spun down at 13,000 rpm for 10 min. The supernatants were saved for PPi measurement while the pellets were solubilized with 2%SDS/62 mM Tris-HCl (pH 6.8) overnight for protein quantification. PPi was measured with a PPi fluorogenic sensor from Sigma (MAK168) in accordance with the manufacturer’s instructions. Briefly, 2 μL of each extracted supernatant was added into a 50 μL assay buffer containing the diluted PPi fluorogenic sensor (1:1000) in a black plate. The mixture was incubated in dark for 20-30 min and read by a plate reader (Tecan infinite 200 Pro) based on the required excitation and emission wavelengths. A PPi standard curve was generated to determine PPi concentrations in samples.

## Acknowledgements

We thank members of the Center for Molecular Parasitology at Drexel University College of Medicine for their technical and intellectual support. We are grateful for plasmid construction assistance from Antara Syam and Dr. Swati Dass. We also thank Dr. Sean Prigge for providing the ACP antibody and the PfMev parasite line, Dr. Akhil Vaidya for providing the KAE609 compound, and PlasmoDB (www.PlasmoDB.org) for providing valuable information. This work is funded by NIH grants (R21AI156735 and R01AI184855) to Dr. Hangjun Ke.

## Conflict of Interest

The authors declare no conflict of interest.

## Notes

### Competing Interest Statement

The authors have declared no competing interest.

